# *PolSpec*: polarisation-based detection for spectral classification of optical signals

**DOI:** 10.1101/2024.10.21.619375

**Authors:** H. Liu, S. Kumar, E. Garcia, P.M.W. French

## Abstract

Spectrally resolved imaging is typically realised using bandpass filters, which are inefficient when they reject “out-of-band” photons or using angularly dispersive devices with at least one image dimension requiring scanning to acquire a full hyperspectral dataset and therefore sequential data acquisition, unless cascades of dichroic beamsplitters are employed, for which cost and experimental complexity scale with the number of spectral channels. Here we present a new approach, “*PolSpec*”, to realise rapid and flexible widefield hyperspectral imaging with lower cost and complexity using polarisation optics instead of dielectric coatings or dispersive devices. *PolSpec* utilises Lyot filters that provide continuously varying transmission across the desired spectral range to generate orthogonal “spectral modulation vectors” that can represent specific spectral signatures with significantly lower data volumes than full spectral profiles. We demonstrate single-shot widefield hyperspectral imaging using a polarisation-resolving camera and rapid, electronically reconfigurable, more photon efficient, hyperspectral imaging using a liquid crystal variable retarder.

## Introduction

Spectrally resolved imaging has been implemented with almost every optical imaging modality, from endoscopy to microscopy and tomography, where spectroscopic information can provide molecular contrast, which is often combined with spatial information in images to identify/classify different states of a sample. It has a wide range of applications including cell and tissue pathology for clinical diagnosis, biochemical assays, spatial proteomics; brightfield and fluorescence imaging of plants, e.g. to assess stress, crop ripeness or the impact of chemical agents (agrochemicals, pollutants), for quality assurance of optoelectronic devices such as solar cells or displays, and other diverse research applications. Multispectral and hyperspectral imaging techniques are also widely used to distinguish and/or unmix light from different chromophores with overlapping spatial distributions in a range of applications. Increasingly, spectrally resolved imaging is used with machine learning to, e.g., increase performance or automate identification/classification processes, e.g., [1,2,3].

Multispectral imaging usually refers to approaches where a relatively small number of discrete spectral channels are resolved and is commonly implemented using a mosaic spectral mask (e.g. RGB camera) or by sequential image acquisitions through either a set of spectral filters or a single electronically tuneable bandpass spectral filter, such as a liquid crystal tuneable filter (LCTF) or an acousto-optic tuneable filter (AOTF). The use of spectral filters is inherently lossy when out-of-band light is rejected and in some implementations of multispectral imaging a (relatively small) number of images are simultaneously acquired using (cascaded) dichroic beamsplitters with multiple detectors - or sometimes a dichroic image splitter is used with a single detector.

Hyperspectral imaging refers to techniques that resolve signals with respect to a continuous set of adjacent spectral channels (bins) so that a “full” spectral profile is acquired for each image pixel. Hyperspectral datasets can also be acquired with sequential image acquisitions via a scanned narrow band filter (typically a LCTF or AOTF) but this is photon inefficient, with the loss increasing with the spectral resolution. For more photon efficient hyperspectral imaging, point scanning or line-scanning instruments can direct signal light to a spectrograph to capture full spectra for the pixels interrogated, in principle, with no out-of-band loss. However, the need to scan along at least one spatial dimension slows down the image acquisition speed.

Although increasingly popular, hyperspectral imaging often acquires more information than is needed for applications such as spectral classification and/or unmixing. Large hyperspectral datasets are often reduced for these purposes using techniques like principal component analysis (PCA) or spectral phasor analysis [4,5,6]. Spectral phasor analysis typically entails multiplying the acquired signal spectra with sine and cosine functions and then integrating them along the spectral dimension to generate vectors on a polar plot (“phasor plot”) that are often referred to as “spectral phasors”. These vectors can represent spectral characteristics, e.g., the mean wavelength and the width of the signals at each image pixel. Where multiple spectral components are present, the spectral phasor is essentially the sum of the component phasors weighted by their respective proportions. If the component (“reference”) spectra are known *a priori*, a spectral phasor can be “unmixed” to give the proportions of each spectral phasor component. In principle, spectral phasors calculated by sine and cosine modulation functions at a single frequency can support linear unmixing up to three spectral components; unmixing more spectral components requires calculation with higher harmonics of the modulation frequency used. Phasor plots can provide convenient visualisations of hyperspectral image data - often enabling different spectral components distributed across a field of view (FOV) to be discerned without further calculations - and this compact (reduced) form of hyperspectral data is amenable to diverse computational and graphical analysis approaches.

Historically, spectral phasor analysis has primarily been applied computationally to “full” (x-y-λ) hyperspectral datasets, which implies that more information was acquired (and more photons detected) than necessary for the intended application. Recently, however, the direct acquisition of spectral phasor data has been demonstrated [ 7, 8 ] by acquiring widefield images transmitted/reflected through filters/dichroics with sinusoidally varying spectral transmission functions. These direct phasor approaches produce much smaller data footprints compared to conventional hyperspectral imaging and provide faster acquisition speeds and/or higher signal-to-noise ratio (SNR) compared to approaches that spread incident photons over many spectral channels. The implementation reported by Dvornikov and Gratton [7] sequentially acquired three transmitted wide-field images: two using sinusoidal and cosinusoidal modulating spectral filters and a third image acquired with no spectral filters, to calculate the spectral phasors across the image. This implementation loses the photons that are not transmitted through the filters, and the sequential acquisitions required mechanical filter changes, limiting the acquisition speed. Subsequently Wang et. al. [8] demonstrated a single-shot implementation, “SHy-Cam”, using two customised dichroics with sinusoidal and cosinusoidal spectral transmission functions respectively. A four-channel image-splitter was used to simultaneously capture 4 widefield images transmitted and reflected from these two special dichroics. This implementation for direct acquisition of spectral phasor data is lossless with respect to the spectral filtering, but the set-up is relatively cumbersome and expensive.

In above demonstrations of direct phasor generation, Dvornikov and Gratton identified off-the-shelf filters for their implementation, while the dichroic beamsplitters used in the SHy-Cam were not standard optical components and so were custom-made. Adjusting the operational parameters of these direct spectral phasor imaging implementations, such as the spectral range, resolution, and higher harmonics of sinusoidal modulation functions would require changing filters and/or dichroic beamsplitters, which may need a new dielectric coating to be designed and fabricated, incurring significant cost. To address the issue of flexibility and to efficiently acquire the information needed for spectral classification and unmixing with reduced cost and minimal experimental complexity, here we propose “*PolSpec*”, where we utilise polarisation optics to implement hyperspectral imaging using “spectral modulation vectors” (SMVs), a generalised approach that includes but is not limited to spectral phasor imaging (where the application of sine/cosine modulation functions is frequently discussed in terms of Fourier transforms). SMV analysis can be used with any convenient sets of continuous orthogonal modulation functions, offering opportunities to reduce experimental complexity.

*PolSpec* utilises common polarisation optics, i.e., linear polarisers, polarising beam splitters, retarders, etc., instead of (bespoke) spectral filters/beamsplitters, to implement spectral modulation functions and thus realise the direct acquisition of SMVs. This approach can widen access to hyperspectral imaging through the wide availability of low-cost polarisation optics, including polymer linear polarising and retarder (quarter and half wave) sheets that can be cut to fabricate components as desired. Electronically configurable flexibility can be implemented using liquid crystal variable retarders (LCRs), e.g., for fast tuning of operational parameters. Here we present two *PolSpec* implementations: one compact, low-cost single-shot hyperspectral imaging module based on a polarisation-resolving (Polarsens™) CMOS camera and an electronically tuneable module based on an LCR combined with an image splitter and a scientific CMOS camera, both of which were implemented on a widefield epifluorescence microscope. As a proof of concept, the instrument response functions (IRFs) of both implementations were measured using a tuneable supercontinuum laser source and the hyperspectral imaging functionality was applied to fluorescence from pollen grain samples.

## Methods

### Generalised Spectral Modulation Vector (SMV)

For an incident spectrum *I*_*i*_(*v*), a *N* -dimensional (normalised) spectral modulation vector (SMV), 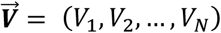, can be calculated by using a set of *N* different spectral modulation functions, {*P*_*k*_(*v*)} where *k* = 1,2, … *N*, and following

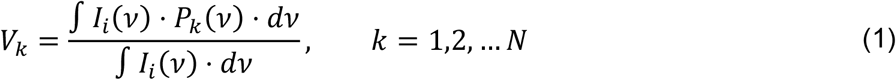

For the convenience of downstream analysis, any two different spectral modulation functions used should be orthogonal, i.e.,

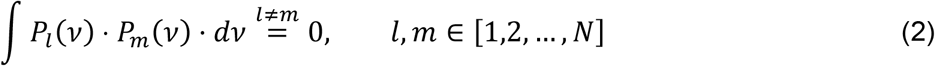

Note that the frequency *v* in Eq.(1)-(2) can also be substituted by the wavelength *λ*, if preferred in practical implementations.

Spectral phasors are specific instances of SMVs generated by {cos(*nfλ*), sin(*nfλ*)} spectral modulation functions, where *f* is the base frequency and *n* is the harmonic number, which is selected as needed in practice. The concept of SMVs extends the choice of spectral modulation functions to *any* sets of continuous orthogonal functions that can be selected to retain the advantages of effective data reduction, e.g., for spectral classification or unmixing applications, while broadening the range of convenient/practical implementations for direct acquisition of SMV image data.

### Polarisation-based Spectral Modulation (*PolSpec*)

For direct acquisition of hyperspectral SMV image data, the required orthogonal spectral modulation functions can be conveniently realised using polarisation optics: *PolSpec* is based on the well-known concept of Lyot filters [9], in which a retarder is usually placed between two parallel (or orthogonal) linear polarisers with its ordinary/extraordinary axes orientated at 45° to the fast/slow axes of the linear polarisers. Figure 1 depicts a configuration for a Lyot filter, whose spectral transmission function *T*(*v*) or *T*(*λ*) for normally incident light is given by the ratio of the output intensity *I*_*o*_ to the input intensity *I*_*i*_ as a function of wavelength (or frequency):

**Figure 1.**
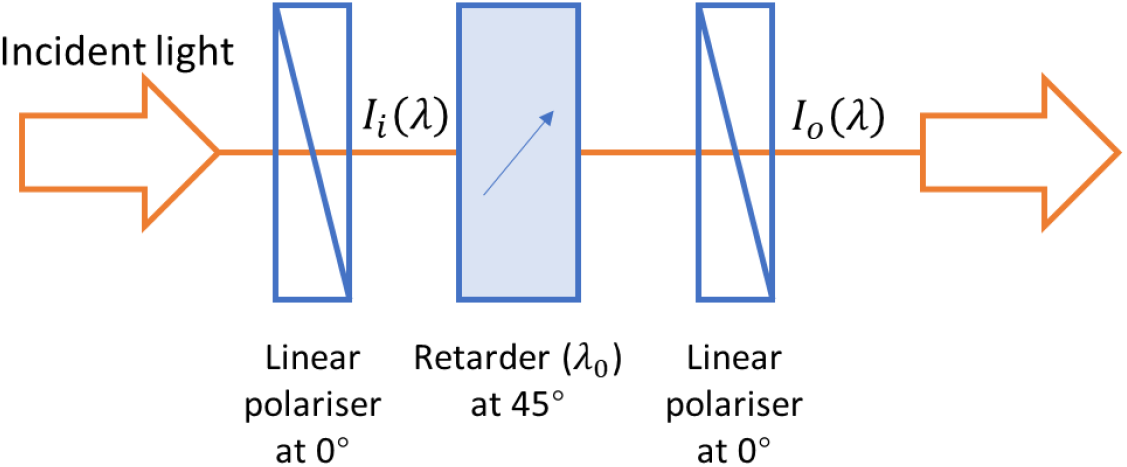
Schematic of a Lyot filter.

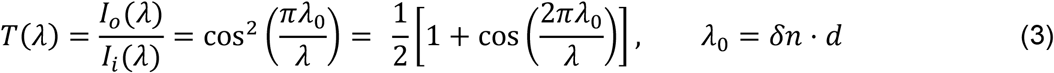

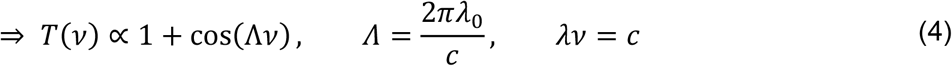

where ∝ represents proportionality, *λ, v*, and *c* denote the wavelength, frequency, and speed of light in free space, respectively. *λ*_0_ denotes the retardance, i.e., optical path difference between the extraordinary and ordinary axes, of the retarder. *δn* denotes the refractive index difference between the extraordinary and ordinary axes of the retarder, and *d* denotes the thickness of the retarder.

From Eq.(4), it is straightforward to configure retarder(s) and linear polarisers to achieve a cosine transmission function. Varying the retardance *λ*_0_ of the retarder(s) results in a change of frequency *Λ* of the cosine function, which consequently changes the spectral modulation function. If spectral phasors are desired, the required sine modulation function requires a frequency-independent quarter wave (π/2) phase shift of the transmission function, which cannot be realised by simply varying the retardance. One possible solution is to insert an achromatic quarter wave plate (AQWP) before or after the retarder. However, some implementations may then involve an adjustment of the optical train and AQWPs are expensive and may not be available for the desired operating spectral range. Therefore, instead of aiming to generate cosine/sine spectral modulation functions for phasor analysis, we instead generate SMVs using a different set of orthogonal spectral modulation functions, {cos(*kΛv*)} where *k* = 1,2, …, which are much easier to configure using polarisation optics and can take advantage of electronic control of the retardance.

### Single-shot *PolSpec* configurations

A single-shot *PolSpec* configuration analogous to the SHy-Cam [8] is shown in Figure 2. The incident light is first divided into two orthogonally linearly polarised beams, *I*_0,*i*_ and *I*_90,*i*_. Each beam then transmits through a retarder with a retardance of *λ*_0_ or 2*λ*_0_, respectively, with the extraordinary/ordinary axes oriented at 45° to the respective beam polarisation. After that, each beam is further split into another two orthogonally linearly polarised beams, both of which are then detected as the complement of each other. Eventually, the readouts from four detectors in Figure 2 are respectively:

**Figure 2.**
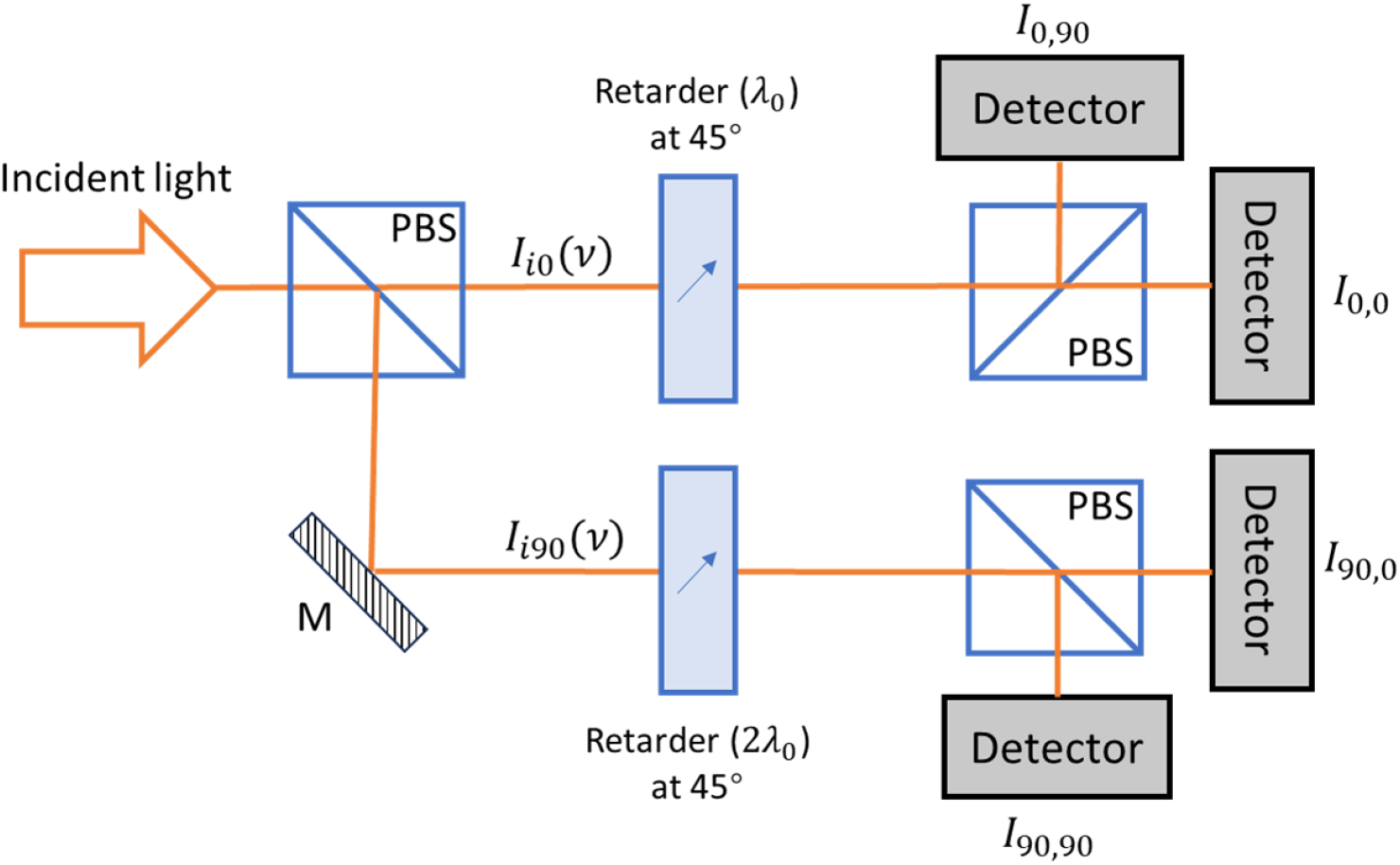
Single-shot PolSpec with {cos(Λv), cos(2Λv)} spectral modulation functions.

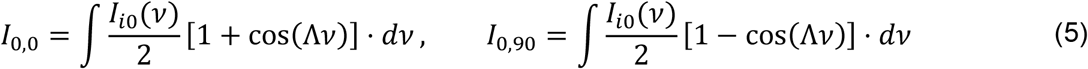

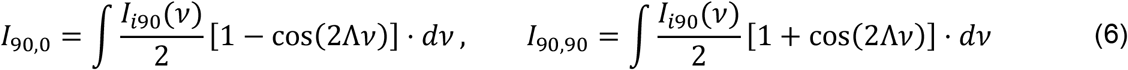

Therefore, 2D SMVs corresponding to the spectral modulation functions {cos(Λ*v*), cos(2Λ*v*)} can be generated by

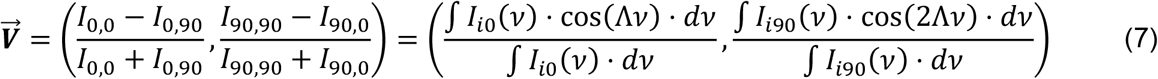

This configuration is single-shot, in principle lossless, and can also provide information about polarisation anisotropy of the incident light. However, like the SHy-Cam [8], it divides the incident beam into four paths, increasing the optomechanical complexity.

A more compact and low-cost configuration of single-shot *PolSpec* can be realised using a polarisation-resolving camera sensor (“Polarsens™” [10]), as shown in Figure 3. A Polarsens™ sensor incorporates a pixel-wise linear polariser mask with fast axes in groups of 2×2 pixels orientated at 0°, 45°, 135°, and 90°, such that a single shot effectively outputs four polarisation-resolved sub-images, *I*_0_, *I*_45_, *I*_135_, and *I*_90_, as if they had been imaged through linear polarisers with fast axes at these orientations.

**Figure 3.**
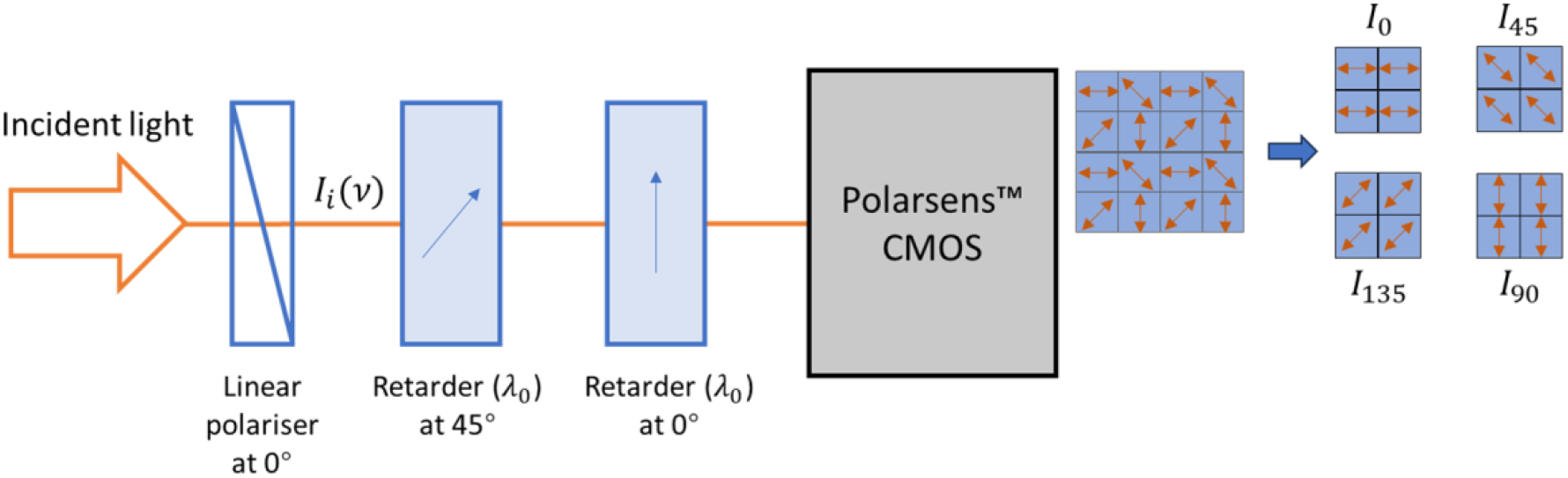
Polarsens™-based single-shot PolSpec with {cos(Λv), sin^2^(Λv)} spectral modulation functions.

The sub-images from the Polarsens™ camera in the set-up depicted in Figure 3 can be respectively written as:

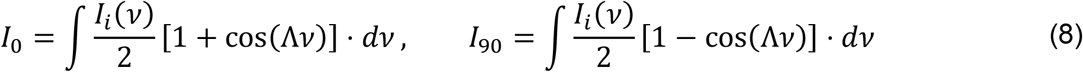

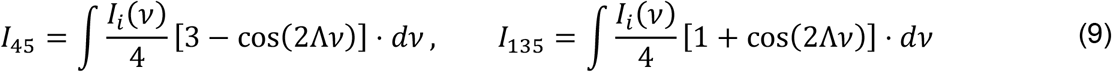

Therefore, 2D SMVs corresponding to the spectral modulation functions {cos(Λ*v*), sin^2^(Λ*v*)} can be generated by:

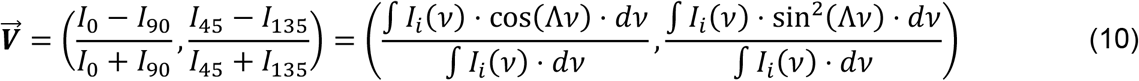

Note that 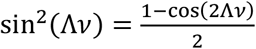 is essentially still a cosine function.

This compact configuration only requires one linear polariser and two retarders, which can be cut from low-cost polymer sheets, and the Polarsens™ camera itself is approximately a cube of side 3 cm weighing 36 g. Such a compact design is suitable for applications with space and/or weight constraints, e.g., on a drone for aerial surveillance or in a handheld device for point-of-care diagnostics. In principle, it may be possible to integrate thin film polarisers and retarders directly in front of the Polarsens™ sensor chip. Supplementary Figure S1 presents an example of brightfield imaging of a colour test chart using the apparatus depicted in Figure 3.

Despite its compact size and low cost, this configuration is limited by the Polarsens™ camera, which is an uncooled CMOS camera designed for machine vision applications. While it enables compact and convenient instrumentation, it presents too much thermal noise for many scientific applications and the pixel-wise polariser mask in front of the Polarsens™ sensor blocks half the light incident, reducing the detection photon efficiency by 50%. This camera exhibits high sensitivity across the visible spectral range, but its polarisation extinction ratio decreases significantly with increasing wavelength from 450 nm, dropping to 100:1 at ∼700 nm [10].

### *PolSpec* configured with LCR

Figure 4 depicts a *PolSpec* configuration that is compatible with a range of imaging modalities, including fluorescence microscopy. An LCR is placed with its (extra)ordinary axis oriented at 45° to the fast axis of input polariser. The output from the LCR comprises two orthogonally linearly polarised components that are detected separately as the complement of each other. By electronically varying the retardance of the LCR to be integral multiples of some *λ*_0_ and acquiring the complementary readouts for each retardance value, *N* dimensional SMVs with the spectral modulation functions, {cos(*k*Λ*v*)} where *k* = 1,2, …, *N*, can be generated. When the retardance of the LCR is *kλ*_0_, two complementary readouts detected can be written as

**Figure 4.**
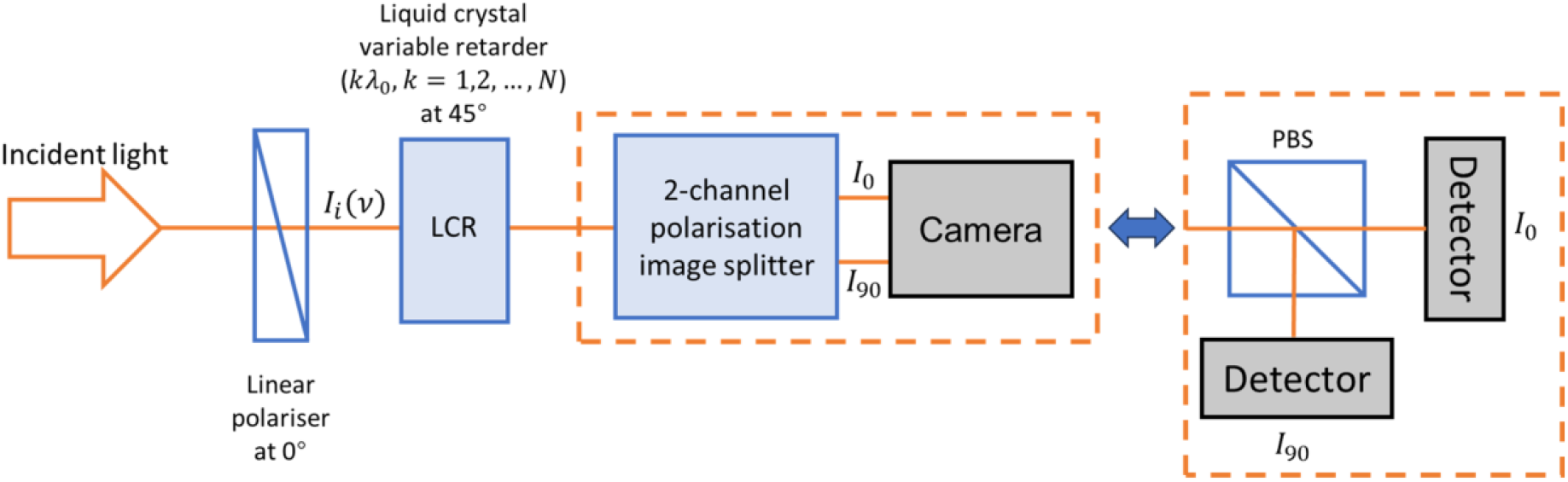
LCR-based PolSpec with {cos(kΛv)}, where k = 1,2, … N, spectral modulation functions.

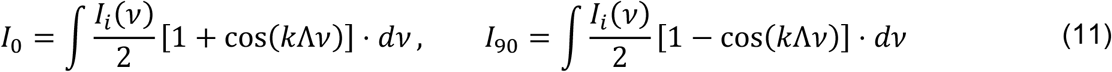

Therefore, the *k*-th elements in the SMVs can be calculated by

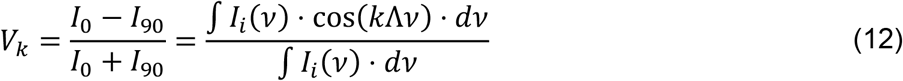

In practical implementations, the value of *N* is usually two but can be increased as needed, e.g., to unmix more spectral components. The upper limit of *N* is determined by the maximum retardance the LCR can reach, although this could be raised by adding additional fixed or variable retarders. With multiple image acquisitions there will be a trade-off with the imaging speed, although we note that LCR can be switched at >100 Hz rates and so *PolSpec* imaging rates of 10’s Hz can be realised.

We note that, compared to the detection in the Polarsens™-based configuration, this is notionally lossless after the first polariser and can be implemented with a wide range of detectors as required by the specific imaging modality. For example, a two-channel polarisation imaging splitter (PIS) combined with a TE-cooled (sCMOS) camera could be implemented for widefield fluorescence. For multiphoton or confocal, scanning microscopy, single pixel detectors such as photomultipliers or line detectors such as CCD arrays could also be incorporated with a polarisation beamsplitter (PBS) for *PolSpec*-based hyperspectral imaging.

### Experimental *PolSpec* implementations

Both *PolSpec* configurations shown in Figure 3 and Figure 4 were implemented as add-on modules to an existing fluorescence microscope frame (Olympus IX71) previously configured for widefield epifluorescence imaging with a tuneable optical fibre laser-based supercontinuum (Fianium, WhiteLase SC400-4) excitation source [11]. The system diagrams of the microscope with the implemented *PolSpec* modules are depicted in Figure 5. In each case, the broadband output from the supercontinuum source was bandpass filtered by a prism-based spectral selection module [11], which provides computer control of the centre wavelength and bandwidth of the excitation light. This excitation beam was homogenized by a rotating diffuser that was imaged to the objective pupil plane by L1 (Thorlabs, AC254-050-A) and L2 (Thorlabs, AC254-300-A) to realise Köhler illumination for epifluorescence microscopy.

**Figure 5.**
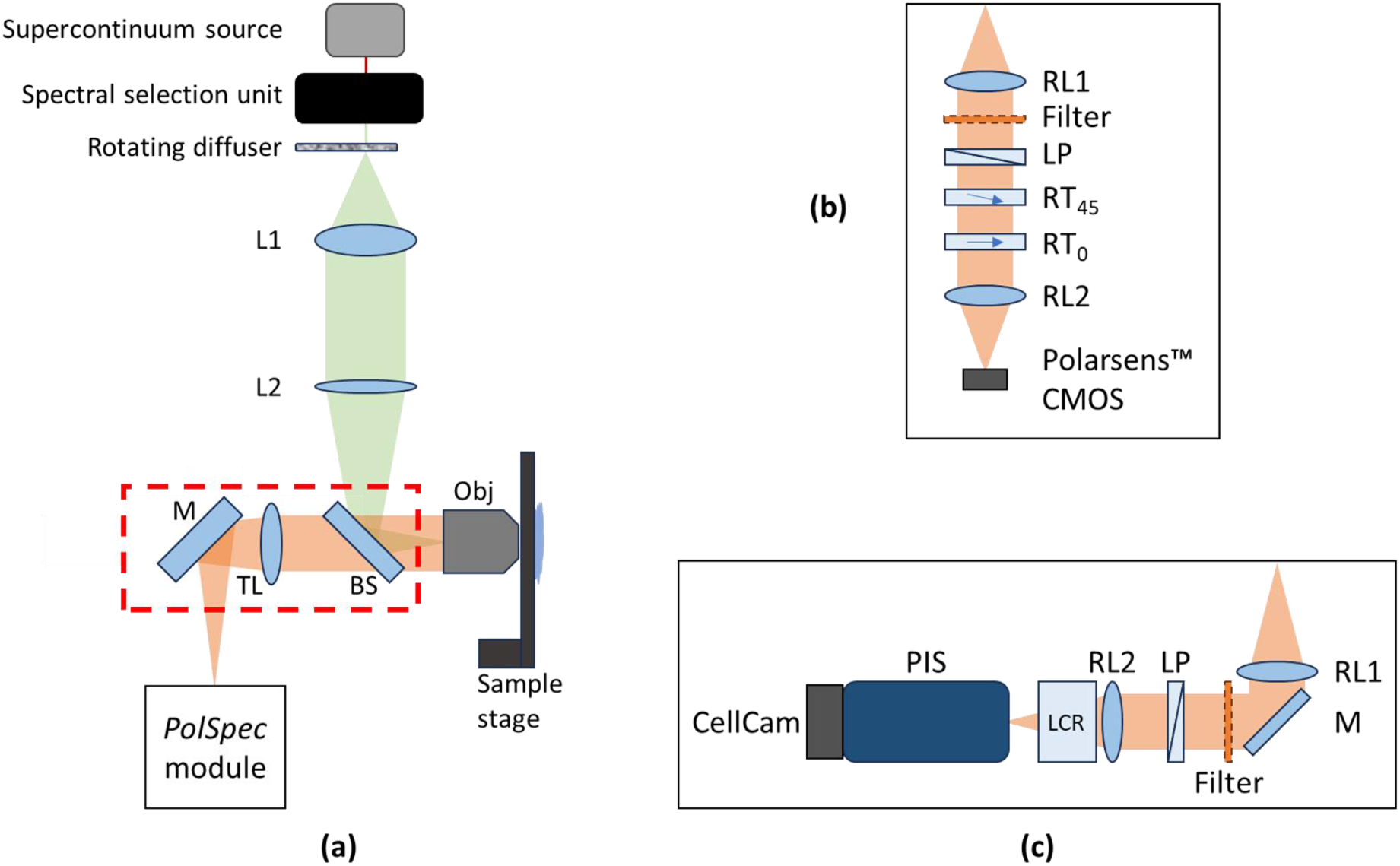
System diagrams of (a) a widefield epifluorescence microscope implemented on an Olympus IX71 frame with (b) a Polarsens™-based single-shot PolSpec module or (c) an LCR-based PolSpec module with a PIS and a sCMOS camera. (L1 & L2, lenses; BS, 50:50 beam splitter; Obj., objective; TL, tube lens; M, mirror; RL1 & RL2, relay lenses; Filter, long pass filter; LP, linear polariser; RT_45_ & RT_0_, retarders with extraordinary axis oriented at 45° & 0° respectively.)

The *PolSpec* modules were mounted at a camera port of the microscope frame, with two identical lenses (Thorlabs, AC254-100-A) relaying the image onto the *PolSpec* detectors. For the Polarsens™-based single-shot configuration, all polarisation optics were placed in the infinity space of this image relay and a Polarsens™ (Teledyne FLIR, Blackfly BFS-U3-51S5P-C) camera was used as the detector. The linear polariser was cut from a low-cost polymer polarising film (Edmund, XP42-40), and the retarders were standard half wave plates designed for 670 nm (Thorlabs, WPH10E-670). For the LCR-based configuration, the same input linear polariser was used and a full-wave LCR (Thorlabs, LCC1423-A) was placed immediately after the second relay lens. A commercial PIS (Optical Insights, Inc, Dual-View™) was mounted in front of a TE-cooled CMOS camera (Cairn Research Ltd, CellCam Kikker 100MT) to serve as the detector.

For the results presented here, the LCR-based implementation was set to acquire 2D SMV image data corresponding to the {cos(Λ*v*), cos(2Λ*v*)} spectral modulation functions in two sequential acquisitions: we set the LCR retardances to ∼700 nm and ∼350 nm respectively, by setting the AC signal applied to the LCR to RMS voltages of 1 V and 1.7 V, at a frequency of 2 kHz. For fluorescence imaging of pollen grains, a long pass coloured glass emission filter (Edmund Optics, Hoya Y52) cutting off at 520 nm was placed before the input linear polariser for both implementations to block the excitation light.

## Results

### Mapping the SMV instrument response functions

The spectral selection unit of the supercontinuum source was set to provide a collimated beam with a spectral bandwidth less than 3 nm and a centre wavelength tuneable between 450 nm and 650 nm. This beam was then directed via a mirror in the sample plane of the microscope to the entrance of the *PolSpec* module, from which the long pass emission filter had been removed. By tuning the centre wavelength of the incident beam, which approximates a spectral delta function, the SMV can be mapped out to provide the IRF of the *PolSpec* module at each wavelength. This is illustrated in Figure 6 where SMVs generated are coloured with their respective wavelength to produce “rainbow response curves” that map the hyperspectral responses of the single-shot Polarsens™-based and (two-shot) LCR-based *PolSpec* modules.

**Figure 6.**
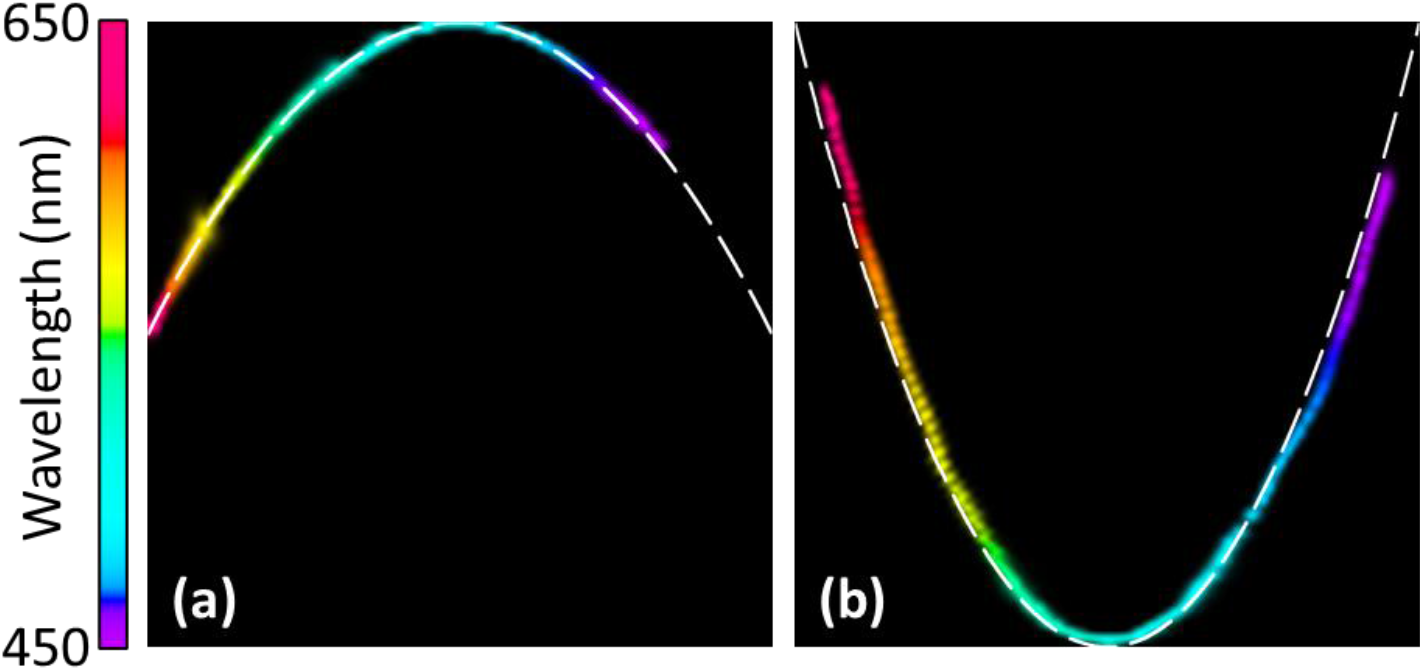
“Rainbow response curves” mapping the hyperspectral IRF of (a) the Polarsens™-based single-shot and (b) the LCR-based two-shot PolSpec modules between 450 nm and 650 nm. White dashed lines indicate the respective theoretical curves for the {cos(Λv), sin^2^(Λv)} and {cos(Λv), cos(2Λv)} spectral modulation functions.

Supplementary video V1 and V2 further demonstrate the cumulative changes in the SMV plot as the centre wavelength of the incident beam was swept from 450 nm to 650 nm. The white dashed lines in the SMV plots indicate the respective theoretical shapes of the response curves. For the Polarsens™-based single-shot implementation, it should be a parabola defined by *y* = 1 − *x*^2^ for the spectral modulation functions {cos(Λ*v*), sin^2^(Λ*v*)}, as sin^2^(Λ*v*) = 1 − cos^2^(Λ*v*). For the LCR-based two-shot implementation, it should be also a parabola, but one defined by *y* = 2*x*^2^ − 1 for the spectral modulation functions {cos(Λ*v*), cos(2Λ*v*)}, as cos(2Λ*v*) = 2 cos^2^(Λ*v*) − 1. The measured response curve for the Polarsens™-based single-shot implementation overlaps well with the theoretical prediction. However, there is small difference between the measured and theoretical curves for the LCR-based two-shot implementation, which could have been caused by the actual retardances of the LCR being different from the theoretical values (which were interpolated from the LCR datasheet), or due to insufficient extinction between two channels of the polarisation image splitter. Nevertheless, the SMVs generated by this *PolSpec* implementation are suitable for application as demonstrated in the following section.

### Spectral classification of fluorescence from pollen grains

With the passband of the spectral selection module set to 423-451 nm and a long pass emission filter installed before the input linear polariser of the *PolSpec* modules, both implementations were applied to image fluorescent pollen grains using a 10x objective (Olympus, UPlanFLN 10x 0.3NA) lens. Figure 7 presents the results for the Polarsens™-based single-shot (top row) and the LCR-based two-shot (bottom row) implementations respectively. The first column in Figure 7 show the total fluorescence intensity images and the second column present the SMV plots generated. Three distinct spectral components can be observed in both SMV plots, from which three masks indicated by the red, green, and blue dashed shaded regions in the orange insets were created. These masks were used to classify every pixel in the intensity image, as shown in the last column of Figure 7, where two groups of pollen grains, coloured by red and green respectively, are differentiated by their fluorescence spectral signatures. The blue component is shown to be background in the fluorescence image, which could be accounted for by the autofluorescence from the mounting media of the pollen grain sample. Note that the un-cooled Polarsens™ camera is not well suited to fluorescence imaging but was able to image the strong fluorescence from this pollen grain sample.

**Figure 7.**
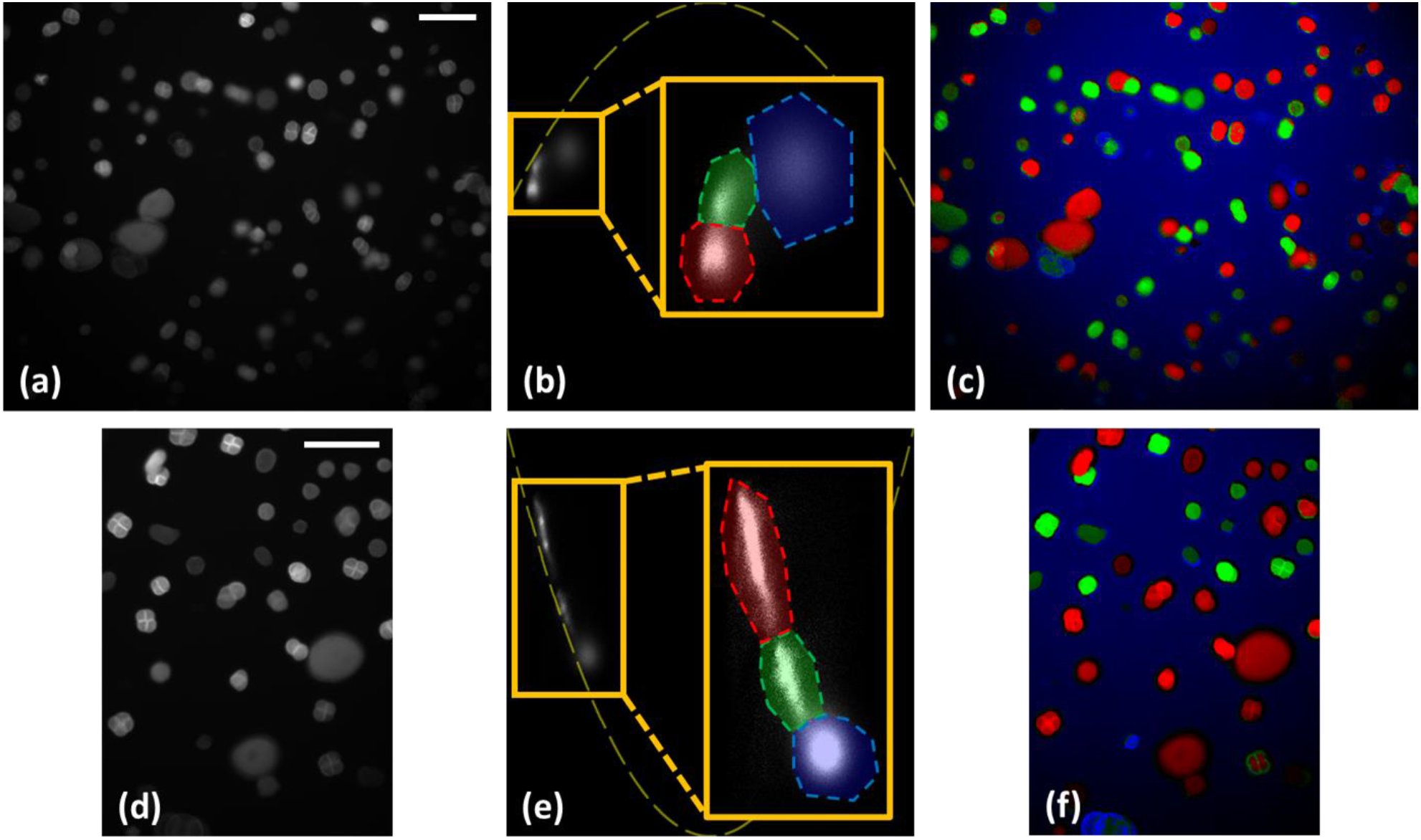
Fluorescence from a pollen grain sample acquired by (a-c) the Polarsens™-based single-shot and (d-f) the LCR-based two-shot PolSpec implementations. From left to right are (a,d) the total fluorescence intensity images, (b,e) the SMV plots with yellow dashed lines indicating the theoretical IRF curves and zoomed insets of the regions of interest used to colour (c,f) the colour-coded total intensity images masked by the dashed shaded regions in the insets of the SMV plots. (Scale bar: 100 µm)

## Discussion

In principle, the spectral range and resolution of *PolSpec* implementations depend on the frequency Λ in the spectral modulation function(s), which is a function of the retardance. Using higher retardances will increase the spectral resolution but decrease the spectral range – although this could be scanned electronically using the LCR. This could be useful for applications requiring high spectral resolution within a limited spectral range, e.g., Raman spectroscopy.

In practice, the spectral discrimination of a *PolSpec* system is limited by the SNR of the detected signals and it is generally desirable to implement *PolSpec* to be as photon efficient as possible. We note that both configurations implemented here require a linear polariser at the input, which causes up to 50% loss for unpolarized incident light such as fluorescence and could be higher for inconveniently polarized incident light. Such input loss could potentially be minimised by placing a wave plate at the input and orientating it to maximize light transmitting the linear polariser. However, it is possible to make *PolSpec* notionally lossless and insensitive to input polarisation by replacing the input linear polariser with a polarising beam splitter (as shown in Figure 2) and implementing two parallel *PolSpec* set-ups for the orthogonally polarised input components. This would also provide information concerning polarisation anisotropy, which could be useful for fluorescence imaging and sensing applications.

Aside from the specific spectral modulation functions, {cos(*k*Λ*v*)} where *k* = 1,2, …, implemented in the work reported here, there are a wide range of other potential spectral modulation functions that can be realised using polarisation optics. As discussed in Methods, it is possible to implement spectral phasors, i.e., the spectral modulation functions {cos(Λ*v*), sin(Λ*v*)}, using *PolSpec* with AQWPs, two possible configurations of which are demonstrated in Supplementary Figure S2 and Figure S3. Another set of *PolSpec* spectral modulation functions, {cos(Λ*v*), sin(Λ*v*) sin(Λ*v*/*X*)} where *X* is a positive integer, could also be realised, as shown in Supplementary Figure S4. By carefully choosing *X* and Λ, the response curve of this configuration can be similar to that of the phasor configuration within certain spectral range, but without requiring AQWPs.

## Conclusions

We have proposed *PolSpec*, a widefield hyperspectral technique based on the direct acquisition of image data to calculate generalized spectral modulation vectors by using polarisation optics to spectrally modulate incident light. *PolSpec* has the same advantages as spectral phasor [7,8] imaging over conventional hyperspectral techniques, including smaller data footprint, faster speed, and better SNR, but may be cheaper to implement and more flexible, particularly if electronically controlled LCRs are utilised. Here two *PolSpec* implementations were demonstrated: a compact single-shot approach based on a polarisation-resolving camera and a tuneable/adaptable approach based on a liquid crystal variable retarder. Their instrument responses were mapped for a spectral range from 450 nm to 650 nm and examples of spectral classification (of fluorescent pollen grains) using both implementations were demonstrated.

*PolSpec* can, in principle, be implemented in almost any optical instrument using any detector where there are polarisation optics available in the appropriate spectral region. This could include wide-field, light-sheet, line-scanning and confocal/multiphoton laser scanning microscopes, endoscopes, telescopes, remote sensing systems and even mobile phone cameras if the polarisation components could be miniaturized. *PolSpec* could be combined with fluorescence lifetime imaging (FLIM) to enable unmixing of more channels using both spectral and lifetime signatures, e.g., for highly multiplexed readouts of fluorescence labels as desired for some spatial proteomics applications. If applied to Förster resonance energy transfer (FRET) measurements, *PolSpec* could enable straightforward recording and unmixing of signals from both donor and acceptor fluorophores for more sophisticated and robust analysis. Some *PolSpec* configurations can also provide polarisation resolved measurements, e.g., to map fluorescence anisotropy.

Potential applications of *PolSpec*-based multispectral and hyperspectral analysis include biomedical and physical sciences research, pathology, in vivo diagnostics (e.g., screening for abnormal tissue potentially indicating cancer), forensics, multiplexed fluorescence readouts (e.g., for sequencing, spatial proteomics, immunofluorescence, etc). It could also be applied to remote sensing for environmental monitoring, surveillance of plants and crops (including from drones), and so on. *PolSpec* data can also be further combined with machine learning to enhance both spectral classification and unmixing applications.

*PolSpec* was presented at the European Microscopy Congress in Copenhagen, August 2024. While preparing this manuscript, we discovered that the use of an LCR to implement hyperspectral imaging in the near infrared using spectral phasors has been reported in arXiv [12], albeit with a (three-shot) less photon efficient implementation than *PolSpec*. This work included an elegant demonstration of the application to image water uptake in plant leaves.

## Acknowledgements

The authors gratefully acknowledge the support of the Optics Instrumentation Facility in the Physics Department at Imperial College London. The practical demonstrations of *PolSpec* have been supported in part by a UKRI Medical Research Council Impact Acceleration award to Imperial College London and by the Chan Zuckerberg Initiative DAF, an advised fund of the Silicon Valley Community Foundation (grant 2021-234618, 2023-321240). Huihui Liu acknowledges a President’s PhD studentship supported by Imperial College London.

## Conflict of Interest

The authors are intending to commercialise aspects of *PolSpec*.

