## Supplementary Information for "*PolSpec*: polarisation-based detection for spectral classification of optical signals"

**A. Hyperspectral imaging of a colour test chart using** Polarsens™**-based single-shot *PolSpec***

**We have also implemented a** Polarsens™**-based single-shot *PolSpec* configuration, following the same setup as Figure 5 (b) except that all polarisation optics used were cut from** cheap off-the-shelf polymer polarising and retarder sheets (**Edmund, XP42-40 & WP280**) at a total cost of less than £100. This implementation was applied to image a colour test chart displayed on a mobile phone screen. Figure S1**(a) provides** the ground truth of colour distribution across the test chart image while Figure S1(b) shows the total intensity image and Figure S1**(c)** shows the SMV plots from three small regions selected from the red, green, and blue areas in the test chart, indicated in Figure S1**(b) using the corresponding colours**. Then, SMVs generated from these regions were plotted in one SMV plot using the respective colours**. As** shown in Figure S1**(c), these three “colours” in the test chart present as three distinct point clouds in the SMV plot, demonstrating its ability to distinguish different spectral components. Three masks were generated from the SMV plot around the point clouds - shown as dashed shaded regions in the inset of** Figure S1**(c). They were then used to “classify” every pixel in the image FOV as red, green, or blue pixels.** Figure S1**(d) demonstrates the intensity image encoded with the classification results, which is consistent with the colour truth in** Figure S1**(a). (Note that pixels not classified to any of three SMV regions were assigned to be black).**

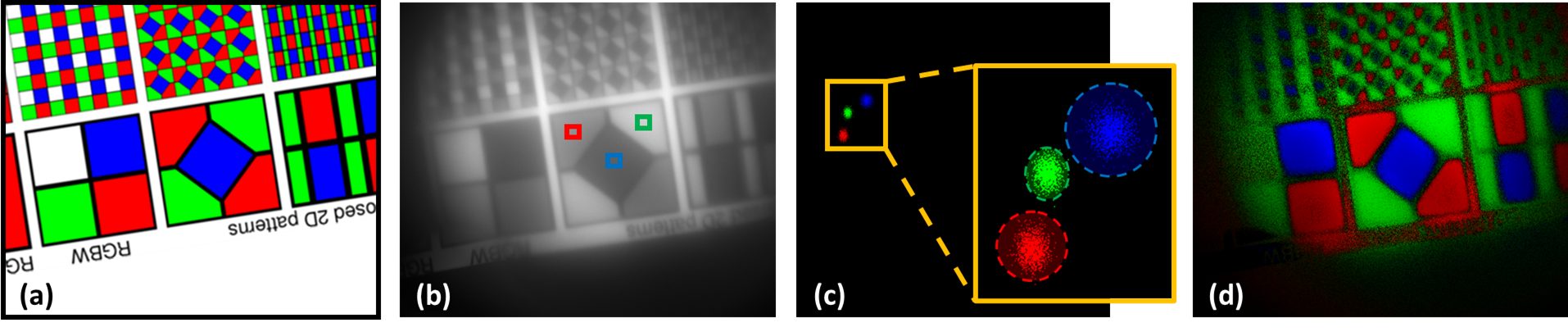

*Figure S1. Colour test chart imaged by a* ***Polarsens™-based single-shot PolSpec implementation:*** *(a) colour test chart ground truth, (b) total intensity image, (c) SMV plot of three bounded regions in (b), (d) colour-coded intensity image generated using masks derived from dashed shaded regions in the inset of (c).*

**B. Potential PolSpec configurations**

Figure S2 and Figure S3 demonstrate two configurations for generating spectral phasors, i.e., SMVs corresponding to the classical spectral modulation functions, $\left\{ \cos\left( \Lambda\nu\right),\sin\left( \Lambda\nu\right) \right\}$, using *PolSpec*. While the configuration in Figure S2 is analogous to SHy-Cam [^[1]^] and quite complicated to implement, the configuration in Figure S3 based on a polarisation-resolving camera (Polarsens™ camera) is more compact. Eq.(1)-Eq.(3) and Eq.(4)-Eq.(6) present the derivations of these spectral phasor modulation functions using polarisation optics for the configurations in Figure S2 and Figure S3, respectively.

For the Figure S2 configuration:

$$I_{0,0}=\int\frac{I_{i0}(\nu)}{2}\left[ 1+\cos\left( \Lambda\nu\right) \right]\cdot d\nu, I_{0,90}=\int\frac{I_{i0}(\nu)}{2}\left[ 1-\cos\left( \Lambda\nu\right) \right]\cdot d\nu$$

(1)

$$I_{90,0}=\int\frac{I_{i90}(\nu)}{2}\left[ 1+\sin\left( \Lambda\nu\right) \right]\cdot d\nu, I_{90,90}=\int\frac{I_{i90}(\nu)}{2}\left[ 1-\sin\left( \Lambda\nu\right) \right]\cdot d\nu$$

(2)

$$\vec{\boldsymbol{V}}=\left( \frac{I_{0,0}-I_{0,90}}{I_{0,0}+I_{0,90}},\frac{I_{90,0}-I_{90,90}}{I_{90,0}+I_{90,90}} \right)=\left( \frac{\int I_{i0}\left( \nu\right)\cdot\cos\left( \Lambda\nu\right)\cdot d\nu}{\int I_{i0}\left( \nu\right)\cdot d\nu},\frac{\int I_{i90}\left( \nu\right)\cdot\sin\left( \Lambda\nu\right)\cdot d\nu}{\int I_{i90}\left( \nu\right)\cdot d\nu} \right)$$

(3)

For the Figure S3*Figure S2* configuration:

$$I_{0}=\int\frac{I_{i}\left( \nu\right)}{2}\left[ 1+\cos\left( \Lambda\nu\right) \right]\cdot d\nu, I_{90}=\int\frac{I_{i}\left( \nu\right)}{2}\left[ 1-\cos\left( \Lambda\nu\right) \right]\cdot d\nu$$

(4)

$$I_{45}=\int\frac{I_{i}\left( \nu\right)}{2}\left[ 1+\sin\left( \Lambda\nu\right) \right]\cdot d\nu, I_{135}=\int\frac{I_{i}\left( \nu\right)}{2}\left[ 1-\sin\left( \Lambda\nu\right) \right]\cdot d\nu$$

(5)

$$\vec{\boldsymbol{V}}=\left( \frac{I_{0}-I_{90}}{I_{0}+I_{90}},\frac{I_{45}-I_{135}}{I_{45}+I_{135}} \right)=\left( \frac{\int I_{i}\left( \nu\right)\cdot\cos\left( \Lambda\nu\right)\cdot d\nu}{\int I_{i}\left( \nu\right)\cdot d\nu},\frac{\int I_{i}\left( \nu\right)\cdot\sin\left( \Lambda\nu\right)\cdot d\nu}{\int I_{i}\left( \nu\right)\cdot d\nu} \right)$$

(6)

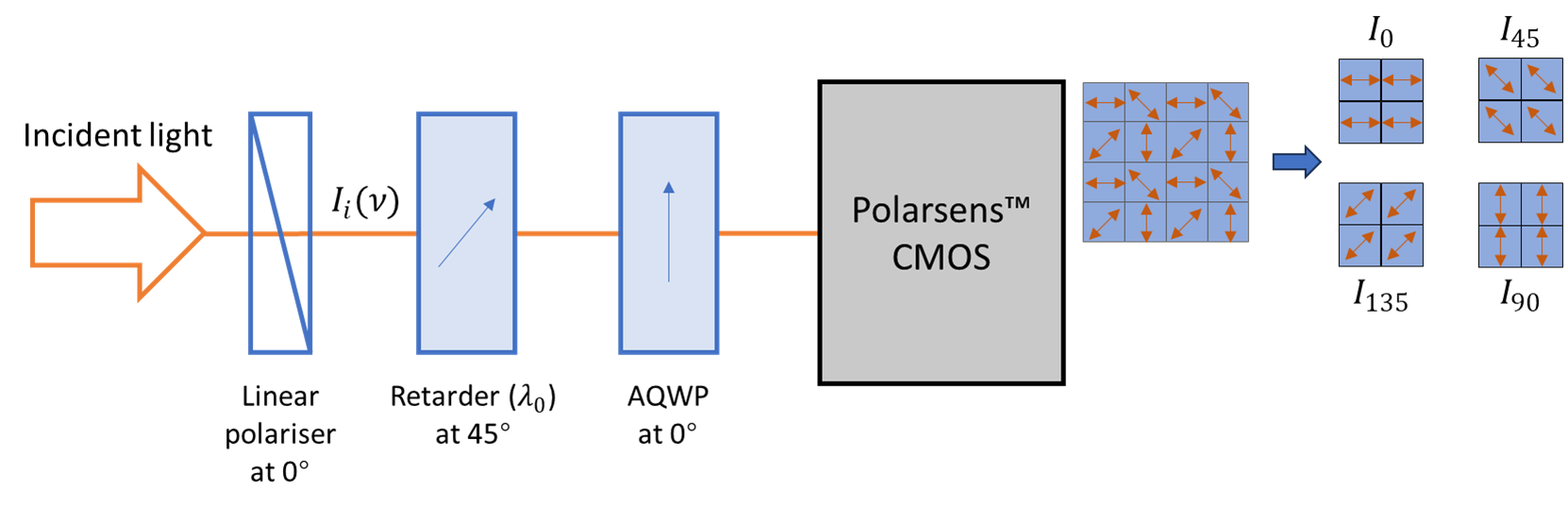

*Figure S3. Polarsens™-based single-shot PolSpec for spectral phasors with* $\left\{ \cos\left( \Lambda\nu\right),\sin\left( \Lambda\nu\right) \right\}$ *spectral modulation functions.*

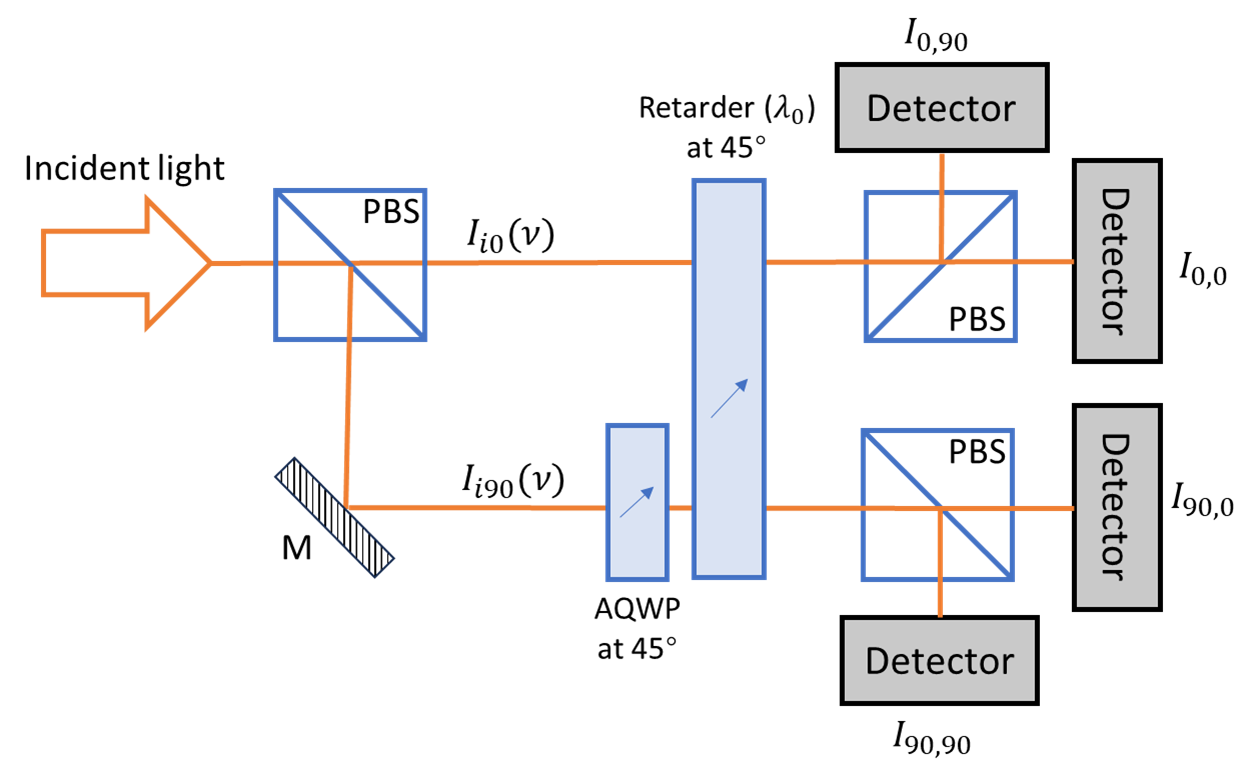

*Figure S2. Single-shot PolSpec for spectral phasors with* $\left\{ \cos\left( \Lambda\nu\right),\sin\left( \Lambda\nu\right) \right\}$ *spectral modulation functions.*

Figure S4 presents a Polarsens™-based single-shot configuration with different spectral modulation functions, $\left\{ \cos\left( \Lambda\nu\right),\sin\left( \Lambda\nu\right)\sin\left( \Lambda\nu/X \right) \right\}$, and the SMV derivation process for this configuration is shown in Eq.(7)-Eq.(9).

$$I_{0}=\int\frac{I_{i}\left( \nu\right)}{2}\left[ 1+\cos\left( \Lambda\nu\right) \right]\cdot d\nu, I_{90}=\int\frac{I_{i}\left( \nu\right)}{2}\left[ 1-\cos\left( \Lambda\nu\right) \right]\cdot d\nu$$

(7)

$$I_{45}=\int\frac{I_{i}\left( \nu\right)}{2}\left[ 1+\sin\left( \Lambda\nu\right)\sin\left( \frac{\Lambda\nu}{X} \right) \right]\cdot d\nu, I_{135}=\int\frac{I_{i}\left( \nu\right)}{2}\left[ 1-\sin\left( \Lambda\nu\right)\sin\left( \frac{\Lambda\nu}{X} \right) \right]\cdot d\nu$$

(8)

$$\vec{\boldsymbol{V}}=\left( \frac{I_{0}-I_{90}}{I_{0}+I_{90}},\frac{I_{45}-I_{135}}{I_{45}+I_{135}} \right)=\left( \frac{\int I_{i}\left( \nu\right)\cdot\cos\left( \Lambda\nu\right)\cdot d\nu}{\int I_{i}\left( \nu\right)\cdot d\nu},\frac{\int I_{i}\left( \nu\right)\cdot\sin\left( \Lambda\nu\right)\cdot\sin\left( \Lambda\nu/X \right)\cdot d\nu}{\int I_{i}\left( \nu\right)\cdot d\nu} \right)$$

(9)

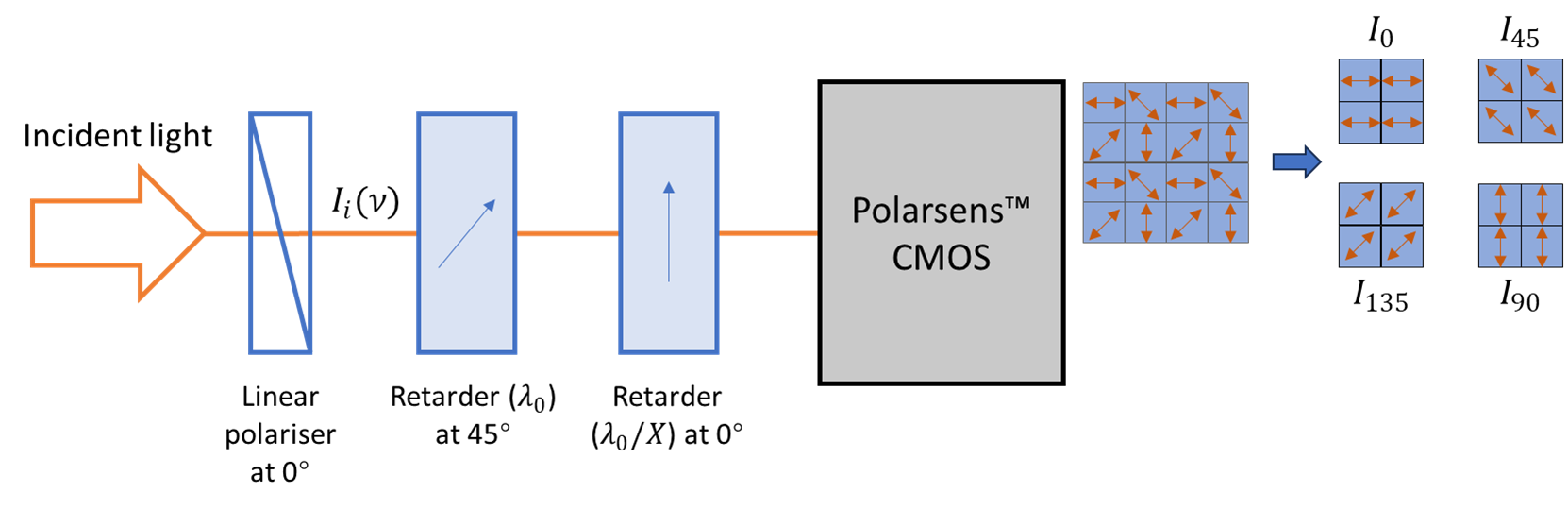

*Figure S4. Polarsens™-based single-shot PolSpec with* $\left\{ \cos\left( \Lambda\nu\right),\sin\left( \Lambda\nu\right)\sin\left( \Lambda\nu/X \right) \right\}$ *spectral modulation functions.*
